# Multiple glacial refugia across northern and southern China and unexpected patterns of spatial genetic diversity in *Betula albosinensis*: a widespread temperate deciduous tree species

**DOI:** 10.1101/2020.10.15.341842

**Authors:** Lu Liu, Andrew V. Gougherty, Junyi Ding, Kun Li, Wenting Wang, Luwei Wang, Feifei Wang, Nian Wang

## Abstract

The central-marginal hypothesis (CMH) predicts high genetic diversity at the species’ geographic centre and low genetic diversity at the species’ geographic margins. However, most studies examining the CMH have neglected potential effect of past climate. Here, we test six hypotheses, representing effects of past climate and contemporary range position, for their ability to explain the spatial patterns of genetic diversity in 37 populations of *Betula albosinensis*. Ecological niche modelling (ENM) revealed large and continuous suitable habitats in north, southwest and southeast China during the last glacial maximum (LGM) but a contraction of suitable habitats since the LGM. Pollen records further confirmed the existence of multiple refugia in north and south China. The spatial pattern of genetic diversity (i.e., expected heterozygosity, gene diversity and allele richness) were best explained by distance to the southern edge and distance to the range edge but also showed longitudinal and latitudinal gradients. Hypotheses accounting for the effects of climate (climatic suitability, climatic stability and climatic variability) had comparatively little support. Our findings show partial support for the CMH and illustrates that the existence of multiple LGM refugia, and suggests species abundance and past species movement play a role in shaping genetic diversity across species’ ranges.

## Introduction

Developing an understanding of the factors shaping the spatial pattern of genetic diversity in species ranges will be important in predicting the response of populations to future climate change and informing population conservation. Populations’ position within a species’ range has been thought to impact the distribution of genetic diversity (Eckert, et al. 2008). For example, one of the most commonly tested hypotheses posits that populations located at the centre of the range tend to have a higher level of genetic diversity than populations at the geographic margin (Eckert, et al. 2008, Micheletti and Storfer 2015). This has been referred as the “central-marginal hypothesis” (CMH), and is thought to result from central populations having higher population abundance, larger effective population size, stronger gene flow and being nearer the ecological optimum compared to marginal populations (Sagarin and Gaines 2002). As a consequence, central populations are expected to have higher genetic diversity within populations and lower genetic differentiation among populations compared with marginal populations (Eckert, et al. 2008, Swaegers, et al. 2013). However, the “central-marginal hypothesis” is often violated as the underlying pattern thought to result in the CMH, i.e. high population abundance in the centre of species’ range is not consistently found (Dallas, et al. 2017, Sagarin and Gaines 2002). Historical climate and species’ response to the changing climate may also impact the spatial pattern of genetic diversity. For example, temperature shifts during the glacial and inter-glacial cycles, especially the last glacial maximum (LGM), often required forest trees to retreat southward into geographically isolated refugia (Hewitt 1999, 2004). As a consequence of this southern retreat and eventual northward recolonisation, many species exhibit a latitudinal decrease in genetic diversity (Petit, et al. 1997); although cryptic refugia and admixture zones are known to obscure simple latitudinal gradients (Petit, et al. 2003). Species’ dispersal ability may also impact the distribution of genetic diversity. For example, species with high dispersal ability sometimes leave a more obvious genetic gradient than species with low dispersal ability (Wang, et al. 2016, Ye, et al. 2019).

The LGM is thought to have had a more minor impact on the distribution of forest trees in East Asia compared to other regions in the northern hemisphere (Qian and Ricklefs 2001, Qiu, et al. 2011). Based on pollen fossil records, it has been proposed that most temperate forest trees in East Asia retreated to between 25 and 30 N during the LGM (Qian and Ricklefs 2001). However, an increasing number of phylogeographic studies indicate the existence of a single refugium in northeast China (NEC) or multiple isolated refugia in NEC and/or north China (NC) for temperate cold-tolerant trees (Hou, et al. 2018, Hu, et al. 2008, Liu, et al. 2012, Zeng, et al. 2015, Zhang, et al. 2005). Several studies indicate an expansion from a single refugium into northeastern China, resulting in a decrease in genetic diversity with increasing latitude (Hu, et al. 2008). Other studies favor multiple isolated refugia in north China, despite a decrease in genetic diversity (Zeng, et al. 2015, Zhang, et al. 2005), or a mixed pattern of genetic diversity, with populations in NEC showing a latitudinal decrease in genetic diversity and a latitudinal increase in genetic diversity in south China (Liu, et al. 2012).

Hence, understanding the drivers of genetic diversity in species ranges requires incorporating information about past climate and demographic history in the frame of CMH. However, to date, only a small number of studies consider past climate (Gougherty, et al. 2020, Jin, et al. 2020). By doing so, some studies have shown that the effects of past climate are relatively minor, while others indicate that historical factors strongly influences the spatial patterns of genetic variation. In addition, many studies do not disentangle historical and contemporary range position and do not account for range-wide spatial auto-correlation.

In this study, we tested whether historical processes affect spatial patterns of genetic diversity in the context of the CMH in China (Wei, et al. 2016). Using *Betula albosinensis* as our study species, we compared six hypotheses for their ability to describe the patterns of genetic diversity across 37 populations. *Betula* is an ecologically important genus that consists of approximately 65 species and subspecies (Ashburner and McAllister 2016, Wang, et al. 2016), widely distributed across the Northern Hemisphere. *Betula* species are wind-pollinated and self-incompatible, often resulting in a high level of genetic diversity within populations (Ashburner and McAllister 2016). Some *Betula* species are pioneers, colonising new habitats and providing shelter for other trees.

Our focal species, *B. albosinensis*, is a deciduous broad-leaved temperate species, growing in mountains at an altitude between 1800~3800m according to our field observations. In China, it extends from northwestern Yunnan province to northern Hebei province, between 25° and 40° latitude and between 99° and 116° longitude. *Betula albosinensis* is a pioneer species and can grow up to 35 meters. Its regeneration depends on habitat disturbances as evidenced by the observations that seedlings are only found in open habitats caused by tree gaps (Guo, et al. 2019).

Here, we used 16 microsatellite markers to investigate the spatial distribution of genetic diversity of *B. albosinensis*. We used ENM and pollen records to robustly infer its suitable habitats since the LGM, and based on these paleo reconstructions of its distribution, we further explored factors impacting the distribution of its population genetic diversity. The specific questions we sought to address are: (1) Does *B. albosinensi*s have multiple LGM refugia in China? (2) Does *B. albosinensis* form different genetic clusters? (3) What is the spatial distribution of genetic diversity? (4) What factors impact the spatial distribution of genetic diversity?

## Materials and Methods

### Field sampling

*Betula albosinensis* samples were collected from natural populations over four years (2016-2019), covering nearly its entire distribution in China. Within each population, samples were chosen at random and separated by at least 20 meters. A GPS system (UniStrong) was used to record sampling locations. Herbarium specimens were collected from most individuals and, for a subset of individuals where twigs were out of our reach, cambial tissue was collected and stored in coffee bags and dried using silica gel. A total of 815 individuals were collected from 37 populations, with 3-56 individuals sampled from each population (Table S1).

### Species distribution model

To predict the potential distribution of *B. albosinensis*, we used occurrence records from our own field work recorded with a GPS system (UniStrong), and collected additional occurrence records from the literature published since the year 2000. If *B. albosinensis* was recorded in a particular certain region, but lacked a geographic coordinate, a random point was selected to represent its occurrence within the region. To avoid spatial autocorrelation due to geographic aggregation, only one point was kept every 5 km. A total of 264 points data were obtained and after filtering, 132 records were retained for ecological niche modelling.

Nineteen bioclimatic variables were downloaded from WorldClim (http://www.worldclim.org) for the four periods: the Last Glacial Maximum (LGM), the Middle Holocene (MID), Present (1970-2000) and Future (2050-2070) (Hijmans, et al. 2005). Present climate variables are derived from monthly mean precipitation and temperature data from the World Meteorological Station from 1970 to 2000 (Fick and Hijmans 2017). Simulated climate data were selected for the other three periods under the Community Climate System Model (CCSM4) (Gent, et al. 2011), in which the data of Representative Concentration Pathways 85 (RCP 85) were selected for FUTURE climate variables. The current and past distribution of *B. albosinensis* were estimated based on an ensemble species model, performed using the R package “BiodiversityR” (Kindt 2018). We selected six bioclimatic variables that lacked strong correlation (<0.75), identified using “ENMTools” (Warren, et al. 2010), to include in subsequent analyses: BIO01 (annual mean temperature), BIO03 (isothermality), BIO07 (temperature annual range), BIO13 (precipitation of wettest month), BIO14 (precipitation of driest month) and BIO15 (precipitation seasonality).

Maximum entropy (MAXENT), generalized boosted regression modeling (GBM), random forest (RF), generalized linear models (GLM) and support vector machines (SVM) were selected for niche model integration simulation test, setting five cross-validations using “BiodiversityR” (Kindt 2018). The five algorithms included in the integrated model had equivalent weights (0.20) for each model. Models were evaluated by splitting data into training and testing datasets with 80% of the data being used to train and 20% to test the models. The true skill statistic (TSS) and the area under the receiver operating characteristics (ROC) curve were used to assess the performance of the models. TSS scores range from −1 to 1, where +1 indicates a perfect ability to distinguish suitable from unsuitable habitat, while values of zero (or less) indicate a performance no better than random. For the ensemble modelling, only those models with a TSS value greater than 0.85 were considered.

### Pollen records

To infer the past distribution of *B. albosinensis* in China, we collected pollen records of *Betula* species from the published literature. For most pollen cores, paleobotanists identified pollen only to the genus level; however, *Betula* pollen in this region is likely *B. albosinensis* as it is currently the dominant *Betula* species. We mapped these pollen sites using coordinates given in the literature. We grouped pollen records into two time periods to coincide with projections from the distribution model: the LGM (22Ka-19Ka) and the Early to Middle Holocene (19Ka-4Ka). The detailed pollen records are listed in table S2.

### DNA extraction and microsatellite genotyping

Genomic DNA was extracted from cambial tissue using a previously modified 2× CTAB (cetyltrimethylammoniumbromide) protocol (Wang, et al. 2013). The quality of genomic DNA was assessed with 1.0 % agarose gels and then was diluted to a concentration of ~10 ng/ul for microsatellite genotyping. Sixteen microsatellite loci developed from closely-related species (Kulju, et al. 2004, Truong, et al. 2005, Tsuda, et al. 2008, Wu, et al. 2002) were used to genotype our samples. The 5’ terminus of the forward primers was labeled with FAM, HEX or TAM fluorescent probes. Each microsatellite locus was amplified individually prior to being artificially combined into four multiplexes. In order to avoid errors caused by size overlapping, loci with significant length differences were labeled using the same dye. PCR procedures were provided in supplementary data. PCR products were delivered to Personal (Shanghai) for microsatellite genotyping. Microsatellite alleles were scored using the software GENEMARKER 2.4.0 (Softgenetics) and checked manually. Individuals with more than three missing loci were excluded, resulting in 815 individuals in the final dataset.

### Microsatellite data analyses

Microsatellite data of *B. albosinensis* was analyzed in STRUCTURE 2.3.4 (Pritchard, et al. 2000) to identify the most likely number of genetic clusters (K) and estimate genetic admixture. As *B. albosinensis* is a tetraploid species, we set ploidy to four in STRUCTURE. Ten replicates of the STRUCTURE analysis were performed with 1,000,000 iterations and a burn-in of 100,000 for each run at each value of K from 1 to 6. We used the admixture model, with an assumption of correlated allele frequencies among populations. Individuals were assigned to clusters based on the highest membership coefficient averaged over the ten independent runs. The number of biologically-meaningful genetic clusters was estimated using the “Evanno test” (Evanno, et al. 2005) and the program Structure Harvester (Earl and vonHoldt 2011). Gene diversity (Nei 1987) and allelic richness (El Mousadik and Petit 1996) were calculated in the software FSTAT 2.9.4 (Goudet 1995). The tetraploid genotypes were treated as two diploid individuals as described by Tsuda et al. (2017) and Hu et al. (2019). A principal coordinate analysis (PCoA) was also performed on microsatellite data of *B. albosinensis* using POLYSAT (Clark and Jasieniuk 2011) implemented in R, ver. 4.0.2 (R Core Team 2020), based on pairwise genetic distances calculated according to Bruvo et al. (2004).

### Geographic and climatic centrality

We used eight variables as predictors of gene diversity (*Gd*), allele richness (*Ar*) and expected heterozygosity (*He*). These include: (a) distance from the geographic range centre (geoCentre), (b) distance from the southern range edge (southernEdge), (c) distance from the range edge (geoEdge), (d) current climatic suitability, (e) climatic distance from the climatic niche centroid (climDist), (f) climatic stability, (g) climatic variability since the LGM, and (h) population structure/genetic admixture. In order to estimate the distribution of *B. albosinensis*, we generated an alpha hull (Rodríguez and Pateiro 2010) around the 264 occurrence records representing the minimum convex polygons (Burgman and Fox 2003) and calculated the distance between each population and the nearest edge. To calculate distance from the geographic range centre, we located the centroid of the alpha hull using “rgeos” (Bivand and Rundel 2018) and then calculated the geographic distance between each population and the centroid (Dallas, et al. 2017, Lira-Noriega and Manthey 2014). The alpha hull was also used to calculate population distance from the southern edge and the distance of each population from the nearest geographic edge (Gougherty, et al. 2020). If populations were located exactly on the geographic boundary or the southern edge, we set the distance to one kilometer.

We calculated the climatic niche centroid by averaging values of the six climatic variables of the 264 coordinate points used for ecological niche modelling. We used Mahalanobis distance as an index to measure the climatic distance between each population and the climatic niche centroid using “statmatch” (D’Orazio 2015). We calculated climatic stability as the sum of suitability of LGM, MID and present (Ortego, et al. 2015, Yannic, et al. 2013); and climatic variability as the standard deviation of suitability of the three time scales (Gougherty, et al. 2020).

In addition to effects of geographic/climatic centrality and past climate, we also tested for effects of population structure and admixture among genetic clusters on *He, Gd* and *Ar*. Using admixture proportions, we calculated a population-level index of admixture. To do so, first we averaged admixture proportions across individuals within populations. We defined populations with genetic admixture between 0.4 and 0.6 as admixed.

We measured the spatial autocorrelation of *He, Gd* and *Ar* using Moran’s Index (I), with −1.0 and 1.0 indicating perfect dispersion and perfect clustering, respectively. Correlograms of Moran’s I for *He, Gd* and *Ar* were estimated in 200 km increments. Significance was determined for both the correlograms and global statistic by comparing the observed statistic to 999 random permutations following Gougherty et al. (2020). Using the eight variables, we compared statistical support for models representing six hypotheses (Table 1). For each hypothesis, we created conditional autoregressive (CAR) models to account for the potential effects of spatial autocorrelation in genetic diversity. CAR models use a weighted estimate of the response variable at neighbouring locations, in addition to the explanatory variables, to parameterise the models (Lichstein, et al. 2002). Neighbourhoods were defined as all populations within 600 km of one another as this distance ensured each population had at least one neighbour and was the approximate maximum distance of continuous positive spatial autocorrelation. Models were compared using Nagelkerke R2, Akaike information criterion (AIC) and Akaike weights (Wagenmakers and Farrell 2004). Each explanatory variable was scaled to a mean of 0 and a standard deviation of 1, to facilitate the comparison of coefficient estimates (Schielzeth 2010).

**Table 1.** Summary statistics for conditional autoregressive models for a range-wide sample of *He*, *Gd* and *Ar* in *Betula albosinensis*, ranked by relative support.

| Model | *He* |  |  | *Gd* |  |  |
| --- | --- | --- | --- | --- | --- | --- |
|  | Nagelkerke<br>R<sup>2</sup> | AIC | AIC<br>weight | Nagelkerke<br>R<sup>2</sup> | AIC | AIC<br>weight |
| southernEdge+GeoEdge | 0.464 | -171.099 | 0.978 | 0.644 | -200.621 | 0.997 |
| southernEdge | 0.289 | -162.655 | 0.014 | 0.479 | -188.459 | 0.002 |
| geoEdge+geoCenter | 0.291 | -160.769 | 0.006 | 0.417 | -182.317 | 0.000 |
| suitability+climDist | 0.184 | -157.946 | 0.001 | 0.253 | -182.793 | 0.000 |
| admixture+maxCluster | 0.227 | -155.560 | 0.000 | 0.443 | -182.008 | 0.000 |
| stability+variability | 0.067 | -155.361 | 0.000 | 0.070 | -178.550 | 0.000 |

## Results

### Ecological niche modelling

A final ensemble model was created by incorporating weighted runs from the MAXENT, GBM, RF, GLM and SVM models, with TSS ranging from □0.89 to 0.92 (TSS average□ = □0.90) and AUC ranging from 0.96 to 0.98 (AUC average = 0.97). The ensemble model predicts high habitat suitability during the LGM in the Qinling-Daba Mountains, northwestern Yunnan and southeastern China (Fig. 1A). During the Holocene, suitable habitat for *B. albosinensis* expanded from the Qinling Mountains into some parts of Shaanxi province and contracted from southeastern China (Fig. 1B). Suitable habitat for *B. albosinensis* seems to have been stable from the Holocene to the present (Fig. 1BC) and is predicted to shift westward in the future (Fig. 1D).

**Figure 1.**
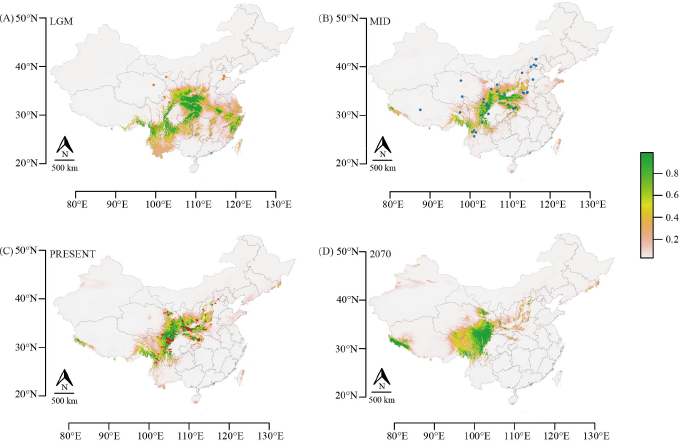
The predicted climatic suitability for *B. albosinensis* and pollen records of *Betula* species. (a) Suitable climate during the LGM, (b) the mid-Holocene, (c) present and (d) in the future (2070). Orange and blue points represent pollen records of *Betula* species during LGM and the Holocene, respectively. Red points represent coordinate points used for ecological niche modelling.

### Pollen records

During the LGM (22 to 19 Ka), eleven pollen records of *Betula* species existed in southwest, central and north China (Fig. 1A). However, pollen records of *Betula* species became more widespread since the beginning of the Holocene (19 to 4 Ka) and appeared in regions representing the current distribution of *B. albosinensis* (Fig. 1B).

### Genetic structure and admixture

Our STRUCTURE analyses identified two clusters (K = 2) as the optimal grouping, according to the ΔK criterion (Fig. S1A). Cluster I, termed hereafter the northern cluster, included populations from the Qinling Mountains and the north (Fig. 2AB). Cluster II, termed the southern cluster hereafter, included several populations from northwestern Yunnan (DQ, JS, LJA, LJS, LP and WX) and southern Sichuan (DC, JD, JLX, LFG, SDX and YJA) (Fig. 2AB). Seven populations (RTX, BMX, MEK, BX, EBY and SLS) located mainly in western Sichuan province showed roughly equal genetic admixture between the northern and southern clusters (Fig. 2AB). The northern and southern clusters were geographically separated by the Sichuan Basin. At higher K values (three to six), populations located at the southern margin in northwest Yunnan provinces (LP, LJS, LJA, JS, WZ and DQ) and on the northwest periphery in Qinghai and Gansu provinces (DTX and XMX) remain genetically distinct (Fig. S1B).

**Figure 2.**
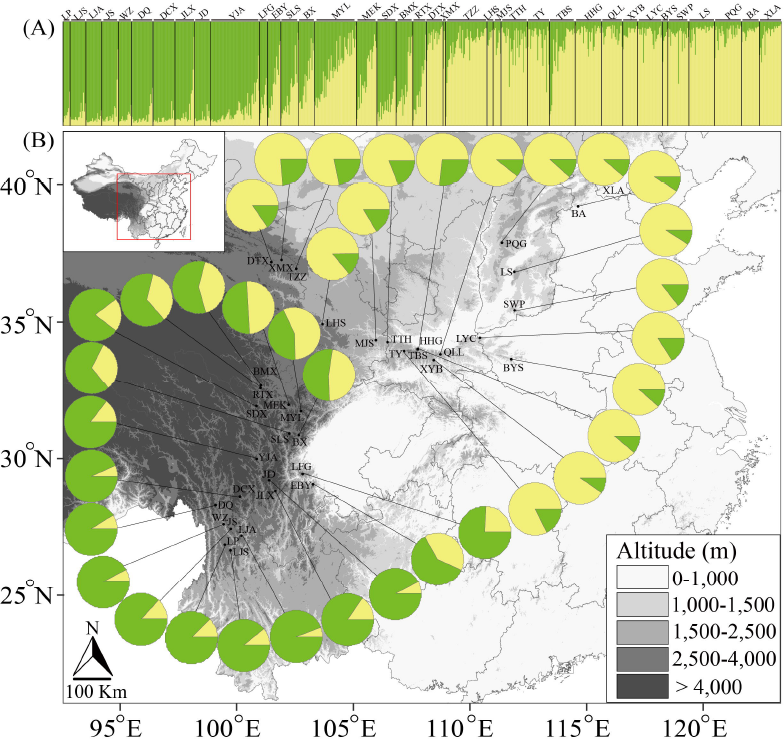
STRUCTURE results for K=2 based on microsatellite markers and a map showing sampling sites. Pie charts indicate the admixture of each population.

We treated individuals with genetic admixture greater than 0.9 from the northern cluster as “northern individuals” and individuals with genetic admixture greater than 0.9 from the southern cluster as “southern individuals”. We treated individuals with genetic admixture in between as admixed. Principal coordinate analysis (PCoA) based on Bruvo’s genetic distances among all samples showed that the “northern individuals” can be separated from the “southern individuals” by PC1 (Fig. 3). Admixed individuals bridged the gap between the “northern individuals” and “southern individuals” and overlapped substantially with individuals from the two clusters. PC1, PC2 and PC3 could not distinguish admixed individuals from individuals from either the northern cluster or the southern cluster. PC1, PC2 and PC3 explained 7.0%, 3.8% and 3.4% of the total variation, respectively (Fig. 3).

**Figure 3.**
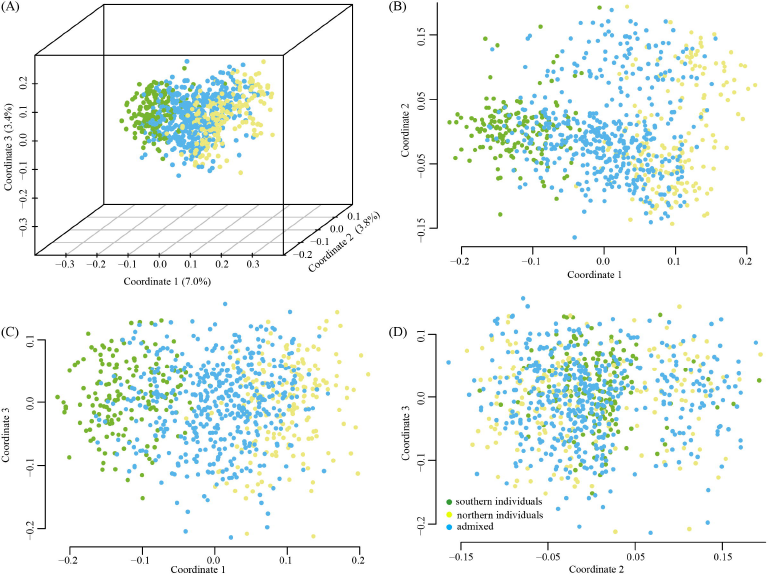
A principal component analysis (PCoA) of *B. albosinensis* at 16 microsatellite markers. Green, yellow and blue points represent “northern individuals”, “southern individuals” and admixed individuals between the northern cluster and the southern cluster, respectively.

### Spatial patterns of genetic diversity

*Gd, Ar* and *He* ranged from 0.72 to 0.83, from 4.04 to 5.07 and from 0.69 to 0.82, respectively. Moran’s *I* showed moderate but significant spatial correlations for *He* (I = 0.19, *P* < 0.001), *Gd* (I = 0.22, *P* < 0.001) and *Ar* (I = 0.32, *P* < 0.001) in the 37 populations, indicating similar levels of genetic diversity among adjacent populations. Correlograms of Moran’s *I* for the three metrics of genetic diversity were positively autocorrelated up to ~200 km, and negatively correlated at 1500 and 2000 km (Fig. 4).

**Figure 4.**
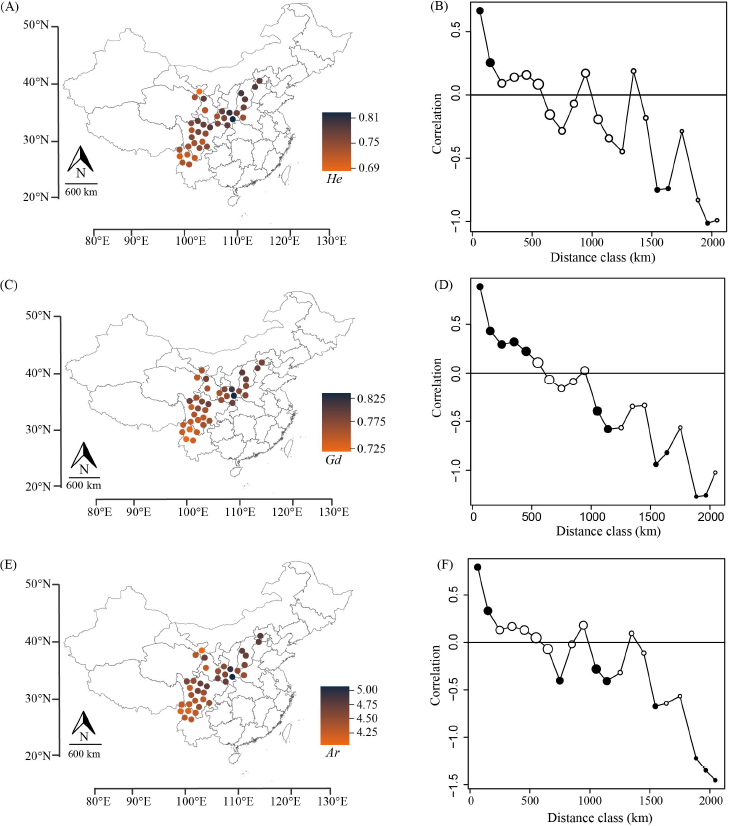
Maps and correlograms (a, b) of genetic diversity (expected heterozygosity = *He*; gene diversity = *Gd*; allele richness = *Ar*) among *B. albosinensis* populations. Circle size in the correlograms is proportional to the number of records within each distance class and filled circles indicate significant autocorrelation at particular distance class (two sided, p > .975 or p < .025).

In general, populations with high *He, Gd* and *Ar* tend to be situated in the northern portion of the range and the geographic centre whereas populations with low *He, Gd* and *Ar* tend to be located near the southern margin. *He, Gd* and *Ar* were positively correlated with both latitude and longitude (Fig. 5). Moreover, we found that climatic distance from the climatic niche centroid was significantly positively correlated (r = 0.49, P < 0.01) with distance from the geographic range centre and significantly negatively correlated with distance from the range edge (r = - 0.64, P < 0.01), indicating that the geographic centre substantially overlapped with the climatic niche centroid.

**Figure 5.**
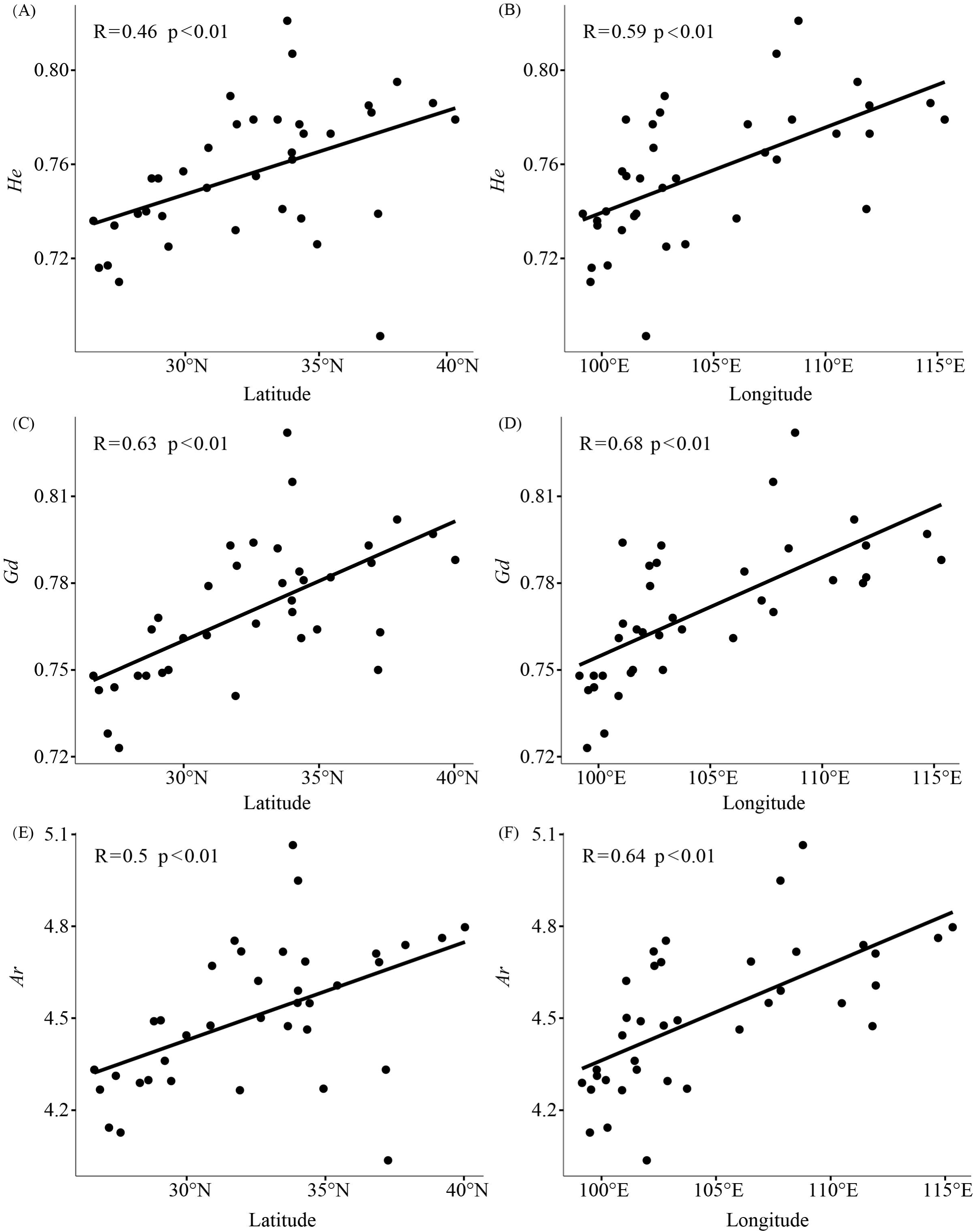
Linear model fit of genetic diversity (expected heterozygosity = *He*; gene diversity = *Gd*; allele richness = *Ar*) with latitude and longitude, respectively.

### Spatial models of genetic diversity

The best performing model for *Gd, He* and *Ar* included distance from the range edge and distance from the southern edge. This model had the highest Akaike weights and highest Nagelkerke R2 (Table 1) for each of the diversity metrics. The spatial models incorporating past climate (climate stability, variability, current suitability and climate distance) tended to have less support, and had Akaike weights near zero (Table 1).

## Discussion

### Range dynamics and its implications

In our study, three lines of independent evidence jointly point towards multiple refugia and cryptic refugia throughout the current distribution of *B. albosinensis*. First, ecological niche models (ENM) predict large and continuous suitable habitats for *B. albosinensis* situated in north, south and southeast China during LGM. Second, the presence of *Betula* pollen dated to the LGM in north and south China further confirms the existence of *Betula* species in these regions. Third, the high genetic diversity within populations in northern China suggests the existence of refugia near these populations. Interestingly, although the ENM does not predict high suitability at LGM at the northern range margin of *B. albosinensis*, high genetic diversity within northern populations (ie., XLA, BA, PQG, LS and SWP) supports the existence of cryptic LGM refugia in this region. This finding, together with several previous studies (Tian, et al. 2009, Zeng, et al. 2015), refutes the hypothesis that temperate forests migrated to the south (25–30 N) during the LGM (Cao, et al. 2014, Harrison, et al. 2001). Instead, our findings favor the “multiple refugia hypothesis” that some temperate trees had multiple refugia in north China (Chen, et al. 2008, Hao, et al. 2018, Wang, et al. 2016).

Unexpectedly, our ENM shows that the suitable habitats for *B. albosinensis* are comparable or even larger than its current distribution, indicating that the LGM may have had little effect on the distribution of *B. albosinensis*. This is possibly due to some unusual characteristics of this species. First, *B. albosinensis* is cold-tolerant and has a broad distribution across north, central and south China. Second, *B. albosinensis* populations occupy a broad altitudinal range between 1800 and 2800m in the Qinling Mountains according to our own field observations. Together, these could indicate that *B. albosinensis* responded to the glacial and interglacial cycles by shifting its range altitudinally, rather than latitudinally. In addition, *B. albosinensis*, like some other *Betula* species, is wind-pollinated and produces a large number of tiny winged seeds (Ashburner and McAllister 2016). This would help *B. albosinensis* disperse long distances and occupy open habitats. Moreover, regeneration of *B. albosinensis* depends on disturbance. Previous studies of *B. albosinensis* in the Qinling Mountains show that its seedlings only grow in open habitats caused by tree gaps (Guo, et al. 2019). These characteristics may have enabled *B. albosinensis* to not only tolerate LGM but may also have allowed it to colonise unglaciated habitats quickly.

Surprisingly, our ENM reveals large suitable habitats for *B. albosinensis* in eastern China during the LGM and disappearance of suitable habitats since the Holocene. *Betula albosinensis* is presently absent from eastern China and *B. luminifera* is the dominant species there. If *B. albosinensis* historically existed in eastern China, one possible scenario for its disappearance is due to its hybridisation with *B. luminifera*, given the extensive hybridisation between *Betula* species documented elsewhere (Anamthawat-Jónsson and Thórsson 2003, Bona, et al. 2018, Eidesen, et al. 2015, Tsuda, et al. 2017, Wang, et al. 2014). Further research characterizing patterns of the genetic admixture between the two species may help to understand this.

### Distinct genetic clusters

Both PCoA and STRUCTURE grouped *B. albosinensis* into two genetic clusters: a northern cluster from the Qinling-Daba Mountains and the regions to the north and northwest, and a southern cluster from northwestern Yunnan and southern Sichuan province. Several populations located in between, such as populations MYL, BX, MEK and EBY showed genetic admixture between the two clusters, indicating regions in between may serve as a contact zone. As expected, the Sichuan Basin is currently a geographic and genetic barrier for *B. albosinensis*, consistent with some previous studies (Qiu, et al. 2009, Wei, et al. 2016). Interestingly, the ENM showed that the Sichuan Basin has been unsuitable for *B. albosinensis* since the LGM, suggesting that it may be partially responsible for genetic differentiation of *B. albosinensis* populations located on both sides of the Sichuan Basin. The absence of *Betula* pollen records from the Sichuan Basin during LGM further indicates the inhospitable habitats for *Betula* species (Fig. 1). Although *Betula* pollen records appeared during the Holocene in Sichuan Basin, it was possibly dispersed from adjacent mountainous ranges, or may belong to *B. luminifera*, which currently has a broad distribution around the Sichuan Basin. In contrast, for *B. albosinensis*, the Qinling-Daba Mountains seems to be a dispersal corridor as populations at both sides of the Qinling-Daba Mountains were not divided into genetically different groups (Fig. 2). Similar results have been observed for other plant species, such as *Fagus sylvatica* (Magri, et al. 2006) and *Fraxinus mandshurica* (Hu, et al. 2008). If mountainous areas are barriers to dispersal, it may depend on several aspects: orientation of mountain range and species dispersal ability (Reeves and Richards 2014). The Taihang Mountains and Lvliang Mountains in north China are of a north-south orientation and the Oinling-Daba Mountains are of a west-east orientation, facilitating the dispersal of *B. albosinensis* along the mountains. Furthermore, the ability of *B. albosinensis* to disperse across the mountains is confirmed by our own observations that it grows on the top of mountains in north China. In general, mountain ranges are more likely to be dispersal barriers for lowland plant species with low dispersal ability. In our case of *B. albosinensis*, its occupation of high altitude and strong dispersal via pollen and seed could allow this species to easily spread across mountains at a broad scale. This explains the low genetic differentiation among populations from north and central China. The northern cluster and the southern cluster of *B. albosinensis* may also reflect local adaptation to different environments, as has been reported for *Quercus aquifolioides* (Du, et al. 2020). It is noteworthy that every individual from the southern cluster has genetic traces from the northern cluster and vice versa, a pattern mirroring incomplete lineage sorting.

### Geographic pattern of genetic diversity

A key finding of our study is that the spatial pattern of genetic diversity of *B. albosinensis* can best be explained by distance from the southern edge and distance from the range edge. Our results partially support the CMH and largely fit the proposed pattern of genetic diversity(Guo 2012). However, the latitudinal effect on genetic diversity is opposite that from most previous studies(Guo 2012). In our study, genetic diversity shows a latitudinal increase whereas many other studies show a latitudinal decrease in genetic diversity. For example, a latitudinal decrease of genetic diversity has been reported for tree species distributed in NEC and subtropical areas due to northward colonisation events (Ye, et al. 2019). Our study also differs with the proposed pattern that genetic diversity decreases from the geographic centre towards both the southern and the northern range (Wei, et al. 2016). Two mutually inclusive hypotheses likely explain a latitudinal increase in genetic diversity: the existence of the northern LGM refugia and a southward movement of *B. albosinensis*. The existence of northern refugia is very likely for *B. albosinensis* as evidenced by ENM results and pollen records and a southward colonisation is also likely due to the dispersal corridor of the Qinling-Daba Mountains.

Interestingly, population HHG, located close to the geographic centre, harbors the highest level of genetic diversity, possibly due to a high abundance of *B. albosinensis*. Based on our field observations, population HHG seemed to be much larger than other populations and had numerous stands of *B. albosinensis*. Hence, we suggest that population abundance may partly account for genetic diversity for *B. albosinensis* in the Qinling Mountains. Although the ENM and pollen records indicate that northwestern Yunnan also served as LGM refugia, populations there had low genetic diversity. Two hypotheses may explain such a pattern: isolated small populations with low species abundance and a southward or an eastward movement of *B. albosinensis*. Unlike populations from the Qinling Mountains where *B. albosinensis* is the dominant species and formed very large and near pure stands in the HHG population, *B. albosinensis* from northwestern Yunnan grows sparsely among other tree species. This may result in the reduced genetic diversity due to genetic drift. Another possibility is that a southward or eastward movement of *B. albosinensis* from the geographic centre or southeastern Tibet, respectively, resulting in loss of genetic diversity due to bottleneck effects. Further inclusion of some populations from Tibet may help to clarify these points. The genetic diversity of *B. albosinensis* is also positively correlated with longitude, with higher levels of genetic diversity residing in more eastern populations. This may indicate a westward dispersal of *B. albosinensis* from central China along the Qinling-Daba Mountains.

Our work also shows that past climate by itself does not explain the patterns of genetic diversity of *B. albosinensis*. For example, the model with past climate stability and variability and model with current climate suitability and climatic distance had low support, with AIC weights being zero and 0.001, respectively. This is somewhat counterintuitive given that *B. albosinensis* seems a demanding species for moisture and temperature. This is possibly due to the fact that the geographic centre of *B. albosinensis* coincides with its climatic centre whereas its peak climatic suitability is not in the geographic centre of the range but in other regions, in particular the western Sichuan and northwestern Yunnan province. Another plausible explanation is that distribution of suitable habitats for *B. albosinensis* did not vary much since the LGM as evidenced by our ENM results. In conclusion, genetic diversity could be related to abundance whereas suitability from ENMs tends not to be associated with patterns of abundance (Dallas and Hastings 2018), resulting in a mismatch between genetic diversity and climatic suitability.

## Acknowledgement

This work was funded by the National Natural Science Foundation of China (31600295 and 31770230) and Funds of Shandong ‘Double Tops’ Program (SYL2017XTTD13).

## Author contributions

NW and AD conceived the project. LL, LW, FW and NW collected samples. LL carried out lab work. LL and AD analyzed the results. NW and AD edited the draft.

**Figures S1** The optimal number of clusters inferred using the “Evanno test” method (A) and (B) STRUCTURE results at K values between 2 and 6 based on microsatellite markers.

**Table S1** Detailed information of populations used in the present study.

**Table S2** Detailed information of pollen records used in the present study.

